# IFITMs Inhibit Cell Fusion Mediated by Trophoblast Syncytins

**DOI:** 10.1101/713032

**Authors:** Ashley Zani, Lizhi Zhang, Adam Kenney, Temet M. McMichael, Jesse J. Kwiek, Shan-Lu Liu, Jacob S. Yount

## Abstract

Type I interferon (IFN) induced by virus infections during pregnancy causes placental damage, though the mechanisms and identities of IFN-stimulated genes that are involved remain under investigation. The IFN-induced transmembrane proteins (IFITMs) inhibit virus infections by preventing virus membrane fusion with cells and by inhibiting fusion of infected cells (syncytialization). Fusion of placental trophoblasts via expression of endogenous retroviral fusogens known as Syncytins forms the syncytiotrophoblast, a multinucleated cell structure essential for fetal development. We found that IFN blocks fusion of BeWo human placental trophoblasts. Stably-expressed IFITMs 1, 2, and 3 also blocked fusion of these trophoblasts, while making them more resistant to virus infections. Conversely, stable knockdown of IFITMs in BeWo trophoblasts increased their spontaneous fusion and allowed fusion in the presence of IFN, while also making the cells more susceptible to virus infection. Overall, our data demonstrate that IFITMs are anti-viral and anti-fusogenic in trophoblasts.

## Main Text

Infections during pregnancy are associated with low birth weights, congenital birth defects, pregnancy complications, and miscarriages (1). Placental damage has been observed during Zika virus infections in non-human primates (2), and in mice during Zika virus and other flavivirus infections (3). Further, a recent landmark study demonstrated that fetal mortality during Zika virus infection is mediated by type I interferon (IFN) receptor signaling(4). This paradoxical effect of what are considered to be beneficial antiviral cytokines may be an evolutionary adaptation in which IFNs signal for termination of pregnancies that are unlikely to be viable due to severe or prolonged infection (4, 5). IFN signaling was shown to disrupt placental architecture in mice, leading to fetal hypoxia and demise. Placental defects caused by IFNs included disruption of the syncytiotrophoblast, a multinucleated cell structure that is critical for nutrient and gas exchange between maternal and fetal blood and that is formed by cell-to-cell fusion of trophoblasts. Though IFNs have been known for decades to be embryotoxic molecules (6-9), the mechanism by which they induce placental damage is not fully understood.

Our laboratory has long studied the IFN-induced transmembrane proteins (IFITMs) 1, 2, and 3, which utilize a palmitoylated amphipathic helical domain to block membrane fusion between viruses and host cells through alteration of lipid bilayers (10-16). These proteins are also capable of blocking cell-to-cell fusion, i.e., syncytia formation, of infected cells (11, 12, 17). Given that placental trophoblast fusion is mediated by unique expression of endogenous retroviral fusion proteins known as Syncytin-1 and −2 (18-23), we hypothesized that IFN and specifically the IFITMs, could inhibit trophoblast fusion and may thus underlie the embryotoxicity of IFN. We sought to test this hypothesis using the human BeWo trophoblast cell line, one of the most commonly used model systems for studying Syncytin-mediated trophoblast fusion (21-27). These cells have a low baseline level of spontaneous fusion, but increase production of Syncytins upon stimulation with forskolin, resulting in robust cell-to-cell fusion (21-26).

Similar to most other cell types, BeWo cells and primary trophoblasts have been reported to produce low steady state levels of IFITM mRNAs even in the absence of IFN (28-30). However, IFITM 1, 2, or 3 were not detected by Western blotting in BeWo cells, indicating that baseline protein levels are low (Fig 1A). Upon IFN treatment, IFITMs 1-3, ISG15, and RIG-I were upregulated, demonstrating functional IFN signaling in these cells (Fig 1A). Co-treatment with fusion-inducing forskolin had no effect on levels of the IFN-induced proteins (Fig 1A). In these treated cells, we quantified cell-to-cell fusion and observed that forskolin treatment significantly increased the fusion index of the cells as expected (Fig 1B,C). This fusion was significantly decreased by IFN co-treatment, demonstrating that type I IFN inhibits fusion of BeWo placental trophoblasts concomitant with induction of IFN-induced proteins, such as IFITMs (Fig 1A-C). The overall level of E-cadherin generally decreases in fused BeWo cells, and was thus moderately decreased in forskolin-treated cells as measured by Western blotting, but was not decreased in forskolin/IFN co-treated cells, providing a secondary indicator of fusion inhibition by IFN (Fig 1B).

**Figure 1:**
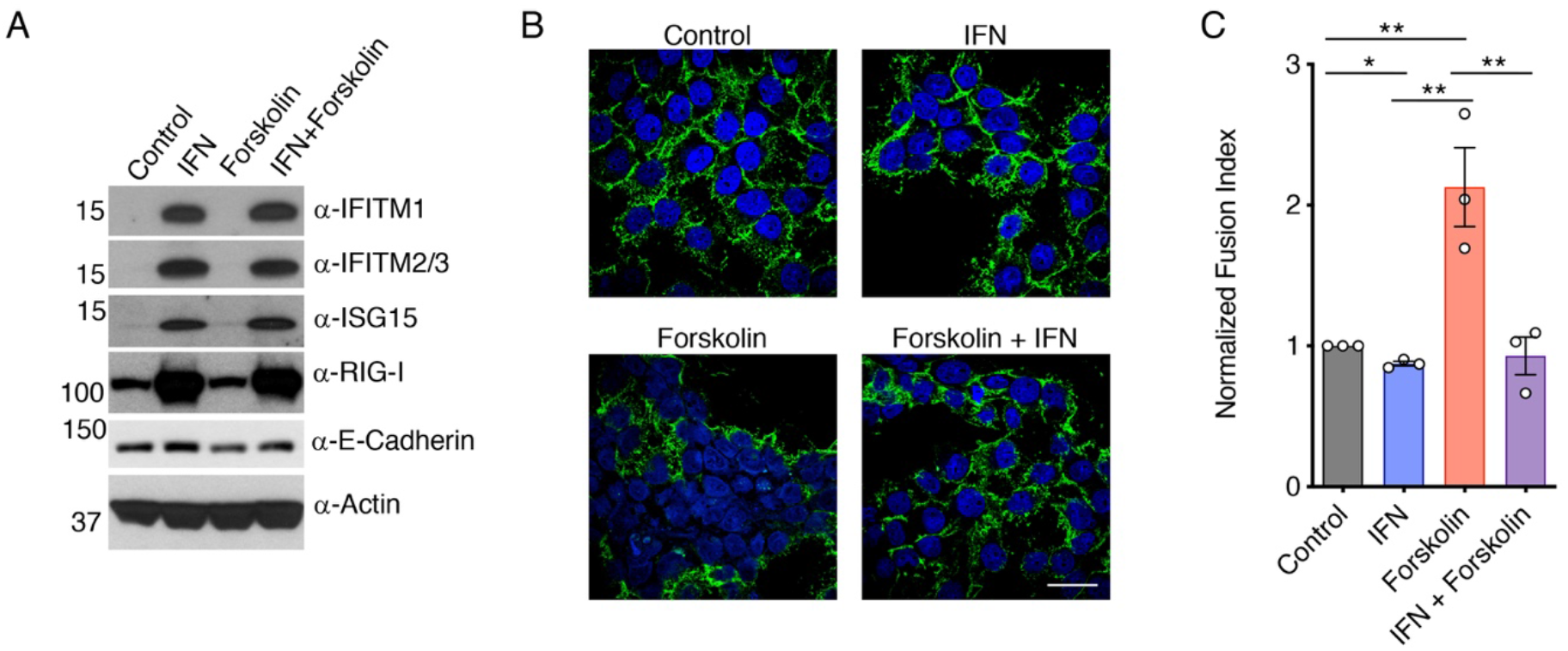
IFN inhibits fusion of BeWo trophoblasts. BeWo cells were treated for 24h with 40 units/mL IFNβ (IFN), 50 uM forskolin, a combination of IFN and forskolin, or vehicle control. Treated cells were analyzed by **A)** Western blotting of cell lysates or **B)** Confocal microscopy imaging after staining with anti-E-Cadherin (green) or DAPI (blue). White scale bar, 10 um. **C)** Fusion indices were calculated and normalized to the background spontaneous fusion level observed in each experiment in vehicle control cells. Bars represent averages from three identical independent experiments with individual data points shown as circles, and data are representative of two additional similar experiments. Error bars represent standard deviation of the mean. *p<0.05, **p<0.01 by paired t-test.

To more specifically examine the effect of the individual IFITMs on trophoblast fusion, we generated stable BeWo lines expressing IFITMs 1, 2, or 3, or an IFITM3 variant (Y20A) that is reported to have decreased endocytic capacity and thus accumulates at the plasma membrane (31, 32) (Fig 2A). To test whether IFITMs are antivirally functional in BeWo cells, we examined infections of the cell lines with influenza A virus and Zika virus. We found that each of the IFITMs decreased infection with influenza virus as compared to vector control cells, and that IFITMs 1 and 3 provided protection against Zika virus infection (Fig 2B). We further observed that forskolin was unable to induce fusion of the IFITM-expressing BeWo lines (Fig 2C,D), indicating that IFITMs 1, 2, and 3 are each capable of inhibiting trophoblast fusion. Thus, IFITM expression in trophoblasts decreases both virus infection and cell-to-cell fusion.

**Figure 2:**
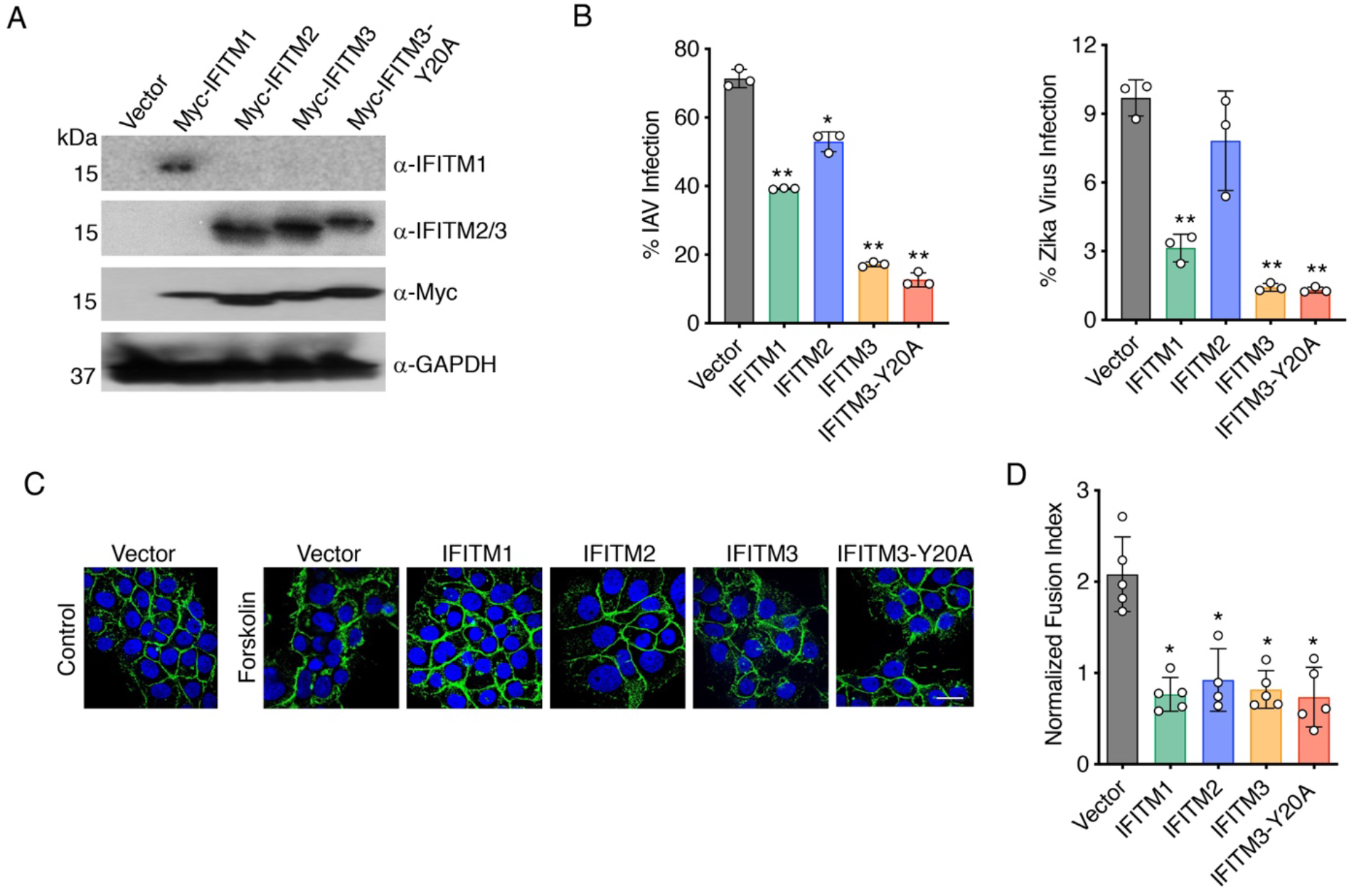
IFITM overexpression inhibits BeWo trophoblast fusion. BeWo cells were stably transduced with lentiviruses expressing myc-tagged IFITM constructs. **A)** Western blotting of cell lysates. **B)** Percent infection 24 h post infection with influenza A virus (IAV) (MOI 2.5) or Zika virus (MOI 5) as measured by flow cytometry staining for viral antigens. Flow cytometry gates were set based on mock infected controls. Bars represent averages of three independent experiments with individual data points shown as circles. Error bars represent standard deviation of the mean. *p<0.05, **p<0.01 as compared to vector control cells by paired t-test. **C)** Cell lines were treated for 48h with 50 uM forskolin or vehicle control. Cells were then imaged by confocal microscopy after staining with anti-E-Cadherin (green) or DAPI (blue). White scale bar, 10 um. **D)** Fusion indices from forskolin-treated cells as in **C** were calculated and normalized to the background spontaneous fusion levels observed in each experiment in vector control cells treated with vehicle control. Bars represent averages from five independent experiments with individual data points shown as circles. Error bars represent standard deviation of the mean. *p<0.05 as compared to forskolin-treated vector control cells by paired t-test.

In order to determine whether IFITMs are specifically capable of inhibiting fusion mediated individually by Syncytin-1 or −2, we employed a cell-to-cell fusion assay utilizing HEK293T cells, which naturally lack expression of endogenous fusion proteins and endogenous IFITMs, and that we previously validated for studying effects of IFITMs on cell-to-cell fusion mediated by viral fusion proteins (12, 17, 33). In short, one population of cells was transfected with vector control or Syncytin-1 or −2, while target cells were transfected with individual IFITM expression plasmids or vector control. Additionally, the two groups of cells were co-transfected with distinct plasmids that produce luciferase only when the plasmids are together in the same cell (34) (Fig 3A). Thus, after mixing the two cell populations, luciferase activity serves as a quantitative readout of cell-to-cell fusion between the two populations (12, 17, 34). We first established that both Syncytin-1 and −2 were able to induce fusion of HEK293T cells (Fig 3B). Fusion mediated by either Syncytin was significantly inhibited by expression of IFITMs in target cells, further establishing that IFITMs are capable of inhibiting cell-to-cell fusion mediated by these essential trophoblast fusogens (Fig 3B). Further, these data demonstrate that inhibition of Syncytin-mediated cell fusion by IFITMs does not require intracellular interaction between Syncytins and IFITMs as these proteins were expressed in distinct cell populations in these experiments.

**Figure 3:**
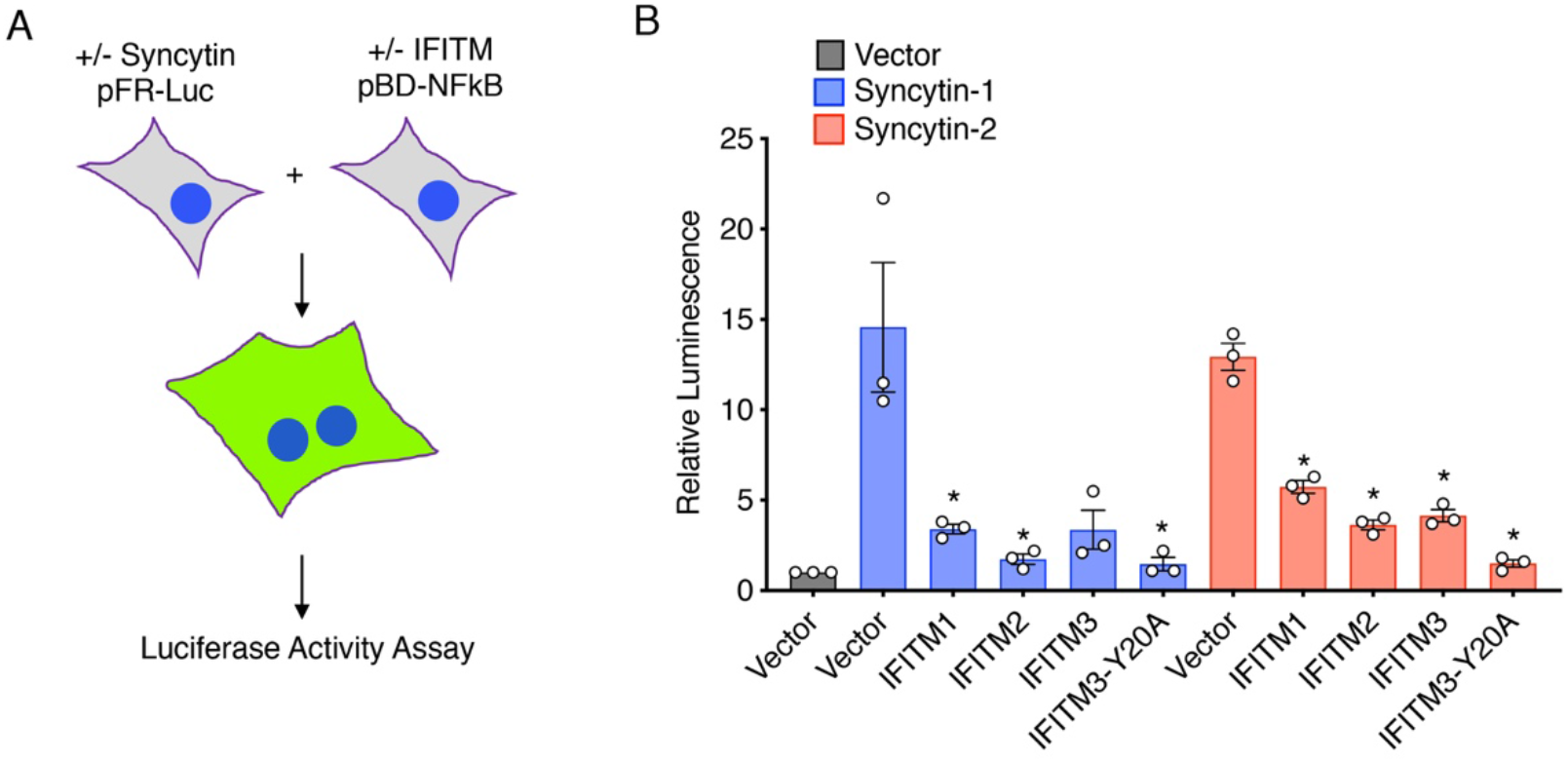
IFITMs inhibition Syncytin-mediated fusion of HEK293T cells. **A)** Schematic representation of the HEK293T cell fusion assay. One population of cells was transfected with Syncytin-1 or −2 constructs or vector control plus pFR-Luc plasmid. A second population of cells was transfected with IFITM constructs or vector control plus pBD-NFkB. Cells were then mixed and allowed to fuse for 24 h. Luciferase activity in cell lysates was then measured as a quantitative readout of cell fusion. **B)** Luciferase activity was measured from cells prepared as described in **A**. Bars represent average results from three independent experiments with individual data points shown as circles. Error bars represent standard deviation of the mean. *p<0.05 as compared to the respective vector controls by paired t-test.

To determine whether endogenous levels of IFITMs are capable of inhibiting trophoblast fusion, we generated stable BeWo cell lines expressing shRNAs targeting the IFITMs. We generated two cell lines (labeled shIFITM #1 and #3) with significant knockdowns of IFITMs 1-3, resulting in low IFITM levels as compared to control cells even with IFN treatment (Fig 4A). Surprisingly, we found that these cells were difficult to maintain when using standard BeWo passaging methods, but that the cells could be readily expanded when passaged frequently at high dilution. We determined that this was likely due to increased spontaneous fusion of these cells and thus their terminal differentiation when grown at high density. We thus quantified fusion of these cells when plated at high density, and found that increased spontaneous fusion was indeed occurring for both of the IFITM knockdown cell lines as compared to control cells, and this fusion was not increased further by forskolin treatment (Fig 4B,C). Strikingly, E-Cadherin was downregulated under these conditions more than we have observed with forskolin treatments, providing a secondary confirmation of the robust spontaneous fusion of these IFITM knockdown BeWo cells (Fig 4D). These results suggest that BeWo cells are capable of fusing without forskolin treatment, but that this is prevented by a low baseline level of IFITMs. Indeed, we were able to detect basal IFITM3 in WT BeWo cells when we prevented IFITM3 turnover in lysosomes by treatment of cells with chloroquine (35) (Fig 4E). Additionally, we observed that IFN was unable to inhibit the spontaneous fusion of these cells, identifying that inhibition of trophoblast fusion by IFN requires the IFITMs (Fig 4F,G). Finally, we measured the Zika virus infection rates of control and IFITM knockdown cells and saw increased infection in the absence of IFITMs (Fig 4H). Overall, we have employed gain- and loss-of-function experiments to identify that IFITMs inhibit virus infection of trophoblasts while also inhibiting critical cell-to-cell fusion required for syncytiotophoblast formation.

**Figure 4:**
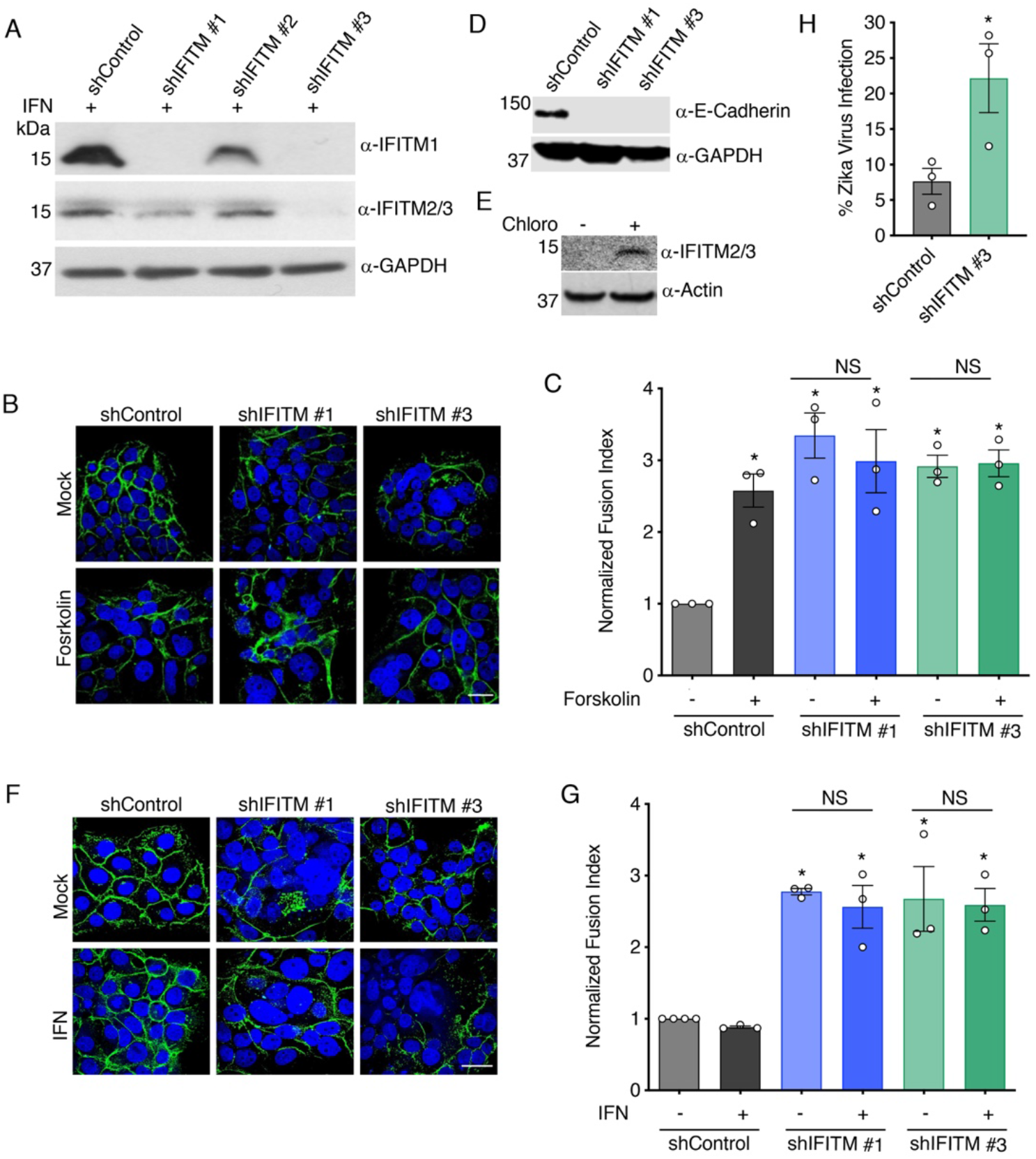
Knockdown of IFITMs promotes fusion of BeWo trophoblasts. BeWo cells were stably transduced with lentiviruses expressing control shRNA (shControl) or different shRNAs targeting IFITMs (shIFITM #1-3). **A)** Western blotting of cell lysates after 18 h treatment with 40 units/mL IFNβ demonstrates IFITM knockdown in specific cell lines. **B)** The indicated cell lines were treated for 48h with 50 uM forskolin or vehicle control. Cells were then imaged by confocal microscopy after staining with anti-E-Cadherin (green) or DAPI (blue). White scale bar, 10 um. **C)** Fusion indices from cells as in **B** were calculated and normalized to the background spontaneous fusion levels observed in each experiment in shControl cells treated with vehicle control. Bars represent averages from three independent experiments with individual data points shown as circles. Error bars represent standard deviation of the mean. *p<0.01 as compared to shControl with vehicle control by paired t-test. NS, not significantly different. **D)** Western blotting of cell lysates demonstrates downregulation of E-Cadherin in IFITM knockdown lines. **E)** Western blotting of WT Bewo cells after 24 h treatment with 40 uM chloroquine (chloro) or vehicle control (mock) demonstrates accumulation of IFITM2/3 when lysosomal degradation is inhibited. **F)** The indicated cell lines were treated for 48h with 40 units/mL IFNβ or vehicle control. Cells were then imaged by confocal microscopy after staining with anti-E-Cadherin (green) or DAPI (blue). White scale bar, 10 um. **G)** Fusion indices from cells as in **F** were calculated and normalized to the background spontaneous fusion levels observed in each experiment in shControl cells treated with vehicle control. Bars represent averages from three independent experiments with individual data points shown as circles. Error bars represent standard deviation of the mean. *p<0.05 as compared to shControl with vehicle control by paired t-test. NS, not significantly different. **H)** Percent infection of shControl cells and shIFITM #3 cells 24 h post infection with Zika virus (MOI 5) as measured by flow cytometry staining for viral antigen. Flow cytometry gates were set based on mock infected controls. Bars represent averages of three independent experiments with individual data points shown as circles. Error bars represent standard deviation of the mean. *p<0.05 as compared to shControl cells by paired t-test.

Together, the results presented here have significant implications for our understanding of the toxic effects of IFNs in infections during pregnancy since they suggest that IFITMs are a critical contributor to this toxicity through effects on placenta syncytiotrophoblast development. The IFITMs may provide antiviral protection during moderate pregnancy-associated infections, including protection against direct viral infections of trophoblasts or the syncytiotrophoblast. However, we posit that IFITMs may also provide a sustained disruption of syncytiotrophoblast formation/maintenance during severe or lengthy infections, resulting in fetal demise. Our data corroborate a groundbreaking recent study showing that IFITMs are required for placental malformations and fetal resorption induced by the viral mimic PolyI:C in mice (27). These observations may also explain the need for tight regulation of baseline cellular levels of IFITM3 by posttranslational modifications, including phosphorylation and ubiquitination, in the absence of infection (14, 31, 35). Interestingly, the antiviral activity of IFITMs is counteracted by the clinically approved drug amphotericin B (17, 36), suggesting that this drug may also hold promise for countering the negative effects of IFITMs on the placenta. Thus, the identification of the IFITMs as a missing link between IFN and placental disruption may provide specific molecular targets for mitigating negative effects of IFNs during pregnancy-associated infections.

## Materials and Methods

### Cell culture and virus infections

An initial stock of Bewo cells was kindly provided by Dr. Stephanie Seveau (The Ohio State University). We confirmed the identity of these cells by confirming their expression of Syncytins and by confirming their fusion in the presence of forskolin. BeWo cells were cultured in Ham’s F-12K media supplemented with 10% Equafetal Bovine Serum (Atlas Biologicals). HEK293T and Vero cells were purchased from the ATCC and were grown in DMEM supplemented with 10% Equafetal Bovine Serum. All cells were cultured in a humidified incubator at 37 °C with 5% CO_2_ and were confirmed to be free of mycoplasma contamination by spot checking with the Lonza MycoAlert Mycoplasma Detection Kit. In some experiments, BeWo cells were treated with IFNβ (EMD Millipore) at a concentration of 40 units/mL. For infections, influenza virus A/PR/8/34 (H1N1) (provided by Dr. Thomas Moran of the Icahn School of Medicine at Mt. Sinai) was propagated in embryonated chicken eggs and titered as previously described (37), and Zika virus strain PRVABC59 (NIH BEI Resources) was propagated and titered in Vero cells. For detection of influenza virus infected cells, anti-nucleoprotein antibody (NIH BEI Resources, NR-19868) staining was used. For detection of Zika virus infection, anti-Zika E antibody (Kerafast) was used. Percent infection was measured by flow cytometry with mock-infected cells serving as controls for gating. Samples were analyzed using FlowJo software.

### Generation of stable BeWo cell lines

For stable expression of IFITM proteins, IFITM coding sequences were cloned into the pLenti-puro vector as described previously (17). Lentiviruses were generated in HEK293T cells and were concentrated using Lenti-X Concentrator reagent (Takara Bio) before transduction of BeWo cells. Stable IFITM-expressing BeWo lines were selected and subsequently maintained with 1 ug/mL puromycin in the media. IFITM knockdown lines were generated and maintained similarly using lentiviral shRNA constructs purchased from Sigma as described previously (38).

### BeWo cell fusion assays

For analysis of cell fusion, BeWo cells were plated to achieve approximately 75% confluence on glass slides. To induce fusion, 50 uM forskolin (Sigma) was added to culture media for 48-72 h. For imaging, cells were fixed with 4% paraformaldehyde, permeabilized with PBS/0.1% Triton-X100, and blocked with PBS/2% Equafetal Bovine Serum. Cells were then stained with anti-E-Cadherin antibody (Abcam, ab1416) at 1:100 in PBS/0.1% Triton-X100 followed by staining with Alexa Fluor 488-labeled goat anti-mouse secondary antibody (Life Technologies) at 1:1000 in PBS/0.1% Triton-X100. Slides were mounted using Prolong Gold Antifade Mountant with DAPI. Cells were imaged on an Olympus FluoView Confocal Microscope. For each sample, 10 randomly chosen images were collected. In each field, nuclei in mono-versus multi-nucleated cells (two or more nuclei within a continuous E-Cadherin stained cell border) were counted. E-Cadherin staining was digitally enhanced using ImageJ software (NIH) to assist with identification of fused and unfused cells. The percentage of fused cells in each field was calculated as nuclei in multinucleated cells divided by the total number of nuclei in the field. Values for each of the ten fields were averaged to determine the percent fusion for each sample. Generally, more than 400 nuclei were counted for each sample. The percent fusion value for the negative control in each experiment (WT BeWo or vector transduced BeWo without forskolin) was set to 1 and all other samples were normalized to this value. The percent spontaneous fusion for the negative controls varied from roughly 10-20% in different experiments and likely represented cells that were fused prior to plating and cells that spontaneously fused during the experiment, and may have potentially also included a small number of false positive multinucleated cells. Fusion induced by forskolin or knockdown of the IFITMs was generally 2-3 fold above the negative control and represented values of roughly 30-80% fusion in different experiments.

### HEK293T cell fusion assays

IFITM constructs cloned into the pCMV-HA vector (Clontech) were described previously (13, 31) and myc-FLAG-tagged Syncytin-1 and 2 expression constructs in the pCMV6 vector were purchased from Origene. pFR-Luc and pBD-NFkB (Agilent) were co-transfected with plasmids as indicated in Fig 3A. Cell-to-cell fusion assays were performed as outlined and validated previously (12, 17, 34). In short, transfections were performed overnight using LipoJet transfection reagent (Signagen). Cell populations were then mixed, replated, and incubated at 37°C in standard culture media for 24 h. Luciferase activity was measured using the Promega Dual-Luciferase Reporter Assay System.

### Western blotting

For Western blotting, cells were lysed in 1% SDS buffer (1% SDS, 150 mM NaCl, 50 mM triethanolamine, pH 7.4), supplemented with cOmplete EDTA-Free Protease Inhibitor Cocktail (Sigma) at 5x the recommended concentration. Western antibodies used in this study include those specific for the following proteins: E-Cadherin (Abcam), ISG15 (Cell Signaling), RIG-I (Cell Signaling), IFITM1 (Cell Signaling), IFITM3 (ProteinTech, detects both IFITM2 and IFITM3), and GAPDH (Life Technologies).

## Acknowledgments

This work was supported by NIH grants AI130110 and AI142256 to JSY. Ashley Zani was supported by NIH training grant AI112542 administered by the OSU Infectious Diseases Institute. The authors thank Dr. Stephanie Seveau for supplying an initial stock of BeWo cells and providing technical advice for fusion assays.

